# Automated extraction and optimization of protein purification protocols using multi-agent large language models

**DOI:** 10.64898/2026.03.03.709341

**Authors:** Jeffery Ye, Amy DeRocher, Monique Khim, Sandhya Subramanian, Lisabeth Cron, Peter J. Myler, Isabelle Q. Phan

## Abstract

Recent advances in Large Language Models (LLMs) present new opportunities for automating critical bottlenecks in scientific workflows such as literature reviews or protocol design. One such bottleneck is the purification of recombinant proteins, a vital aspect of biomedical research that frequently fails. To improve success rates, researchers must manually define optimal large-scale purification conditions and establish robust rescue protocols for proteins with low stability or solubility – a time-intensive process.

To address this gap, we introduce a multi-agent LLM system that automates the creation and optimization of protein purification protocols to facilitate the production of high-concentration, high-purity protein samples. Our application streamlines the labor-intensive manual process of sequence similarity searches, literature reviews, and protocol comparison. Operating in a tool-like constrained workflow, the system identifies analogous proteins, leverages specialized LLM agents to extract successful purification methodologies from primary source literature, and cross-references them against failed protocols to generate optimization recommendations.

Evaluation on a select number of targets demonstrated high accuracy in protocol extraction and the generation of scientifically sound, expert-validated optimization recommendations. While this system reduces complex analysis time from hours to minutes, we identify the lack of programmatic open access to literature, specifically primary citations in the Protein Data Bank, as a fundamental limitation to LLM agent-based scientific workflows. Ultimately, this system demonstrates the feasibility of using LLM agents to streamline wet-lab workflows while preserving methodological transparency and reproducibility.

## Introduction

Research tasks such as literature reviews or experimental protocol design represent critical bottlenecks in scientific workflows, often consuming hours of expert time on highly structured, repetitive analysis (Luo, et al., 2024). Recent advances in Large Language Models (LLMs) present new opportunities for automating these language-intensive tasks with higher accuracy and efficiency than manual approaches (Scherbakov, et al., 2025), (Kang, et al., 2025), (Li, et al., 2024). However, the potential for LLM enhancement of these tasks in a wet lab setting remains largely untapped, partly due to limited experience among researchers in developing AI applications for scientific workflows (Giesriegl, et al., 2025), (Bianchini, et al., 2025). Current applications of LLMs in biomedical research typically focus on generalized AI assistants that may struggle with the specificity and reproducibility required for rigorous scientific workflows (Huang, et al., 2025), (Su, et al., 2024). While these broad-domain LLMs can theoretically perform literature analysis tasks, they fail to provide standardized, transparent methodologies essential for many research applications (Lin, et al., 2024). To address these limitations, this study introduces a specialized LLM multi-agent workflow to accelerate discovery, by reducing a multiple-hour manual workflow to a two-minute automated analysis, while maintaining reproducibility and methodological transparency. Our LLM system serves as a representative application of how multi-agent architectures can benefit biomedical research (Klang, et al., 2025).

High-yield, high-purity, and high-activity protein samples are essential for the success of diverse biomedical applications, including structural biology, proteomics, and drug discovery (Raynal, et al., 2014). At the Seattle Structural Genomics Center for Infectious Disease (SSGCID), we possess a high-throughput gene-to-structure pipeline (Stacy, et al., 2011), attempting over 10,200 purifications across 5,500 distinct targets from over 250 different organisms. However, our success rates have remained consistently low, where barely a third of cloned targets successfully purify. Even when common expression challenges are overcome (e.g. optimized construct design, codon usage, and expression vectors/media) (Rosano, et al., 2014) and soluble expression is successful, the subsequent large-scale purification step still fails in over 30% of cases, a rate that has remained stable over time (internal data).

An established workflow for rescuing failed purifications typically includes searching for similar proteins from related organisms in the Protein Data Bank (PDB), extracting the experimental protocols from associated primary citations, and comparing critical parameters with the failed protocol (Gileadi, et al., 2007). This manual process can take multiple hours per target. The structured, sequential nature of this workflow makes it an ideal candidate for an agentic LLM solution. A multi-agent architecture is particularly well-suited to this task because the rescue workflow involves distinct sequential steps that require accurate and specific expertise. Dividing tasks between multiple models can reduce hallucinations and inaccuracies, allowing each LLM to be an expert at its specific role (Klang, et al., 2025).

Here we introduce a multi-agent LLM system (Figure 1) that automates each step in the workflow. Using custom built taxonomic and sequence similarity tools, our system identifies successful purifications of proteins similar to the rescue target, summarizes the relevant literature, and modifies any failed protocols to generate a comprehensive report on how to purify the troublesome protein. Our findings suggest that LLMs can effectively streamline repetitive research workflows, enabling researchers to prioritize tasks that require human intuition and analytical reasoning.

**Figure 1.**
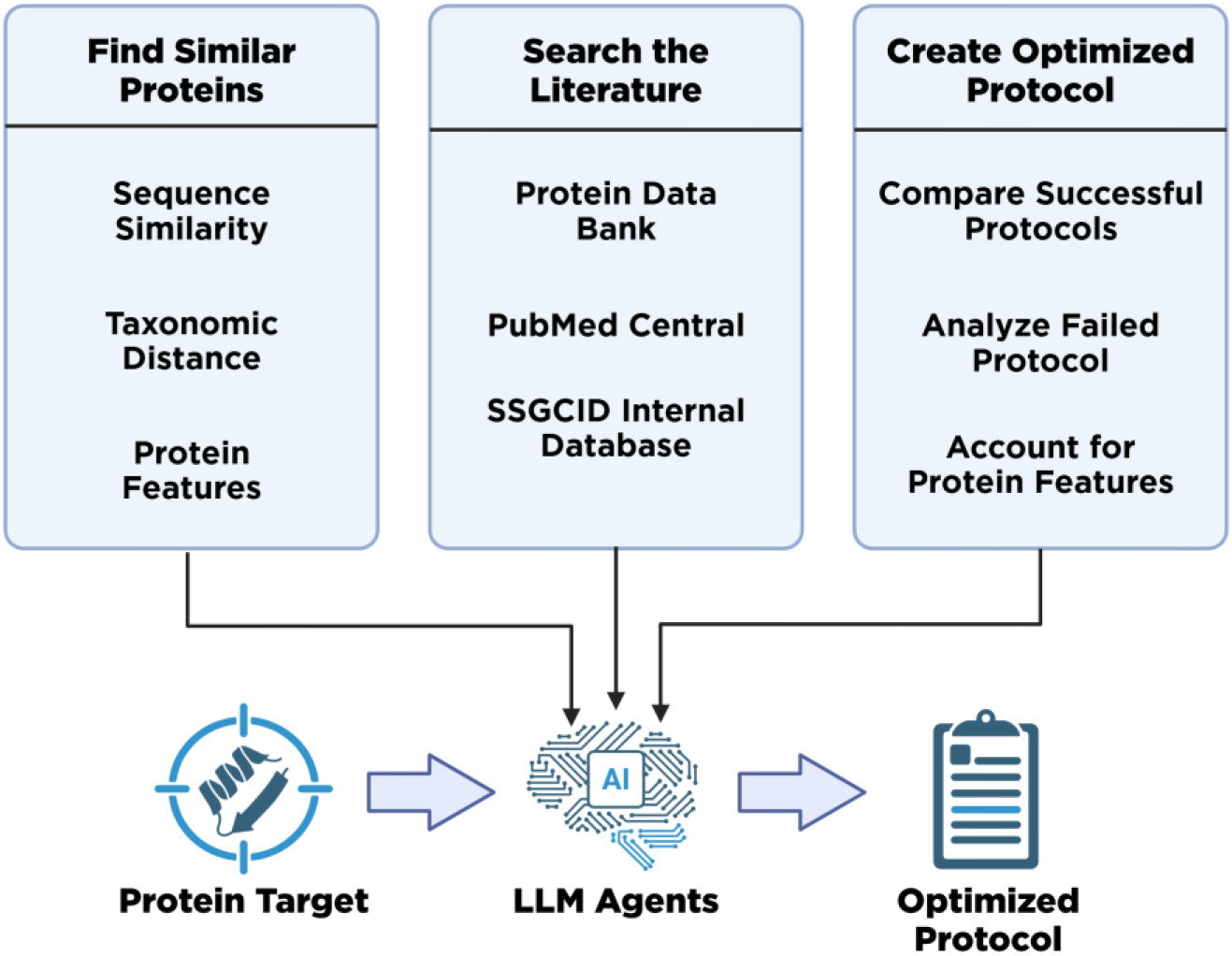
Overview of the multi-agent system integrating bioinformatics tools (BLAST, taxonomy tree navigator), literature sources (PubMed Central, SSGCID Database), and AI agents using native PydanticAI validation to automate protein purification rescue workflows.

## Methods

### Similarity Calculation

Proteins analogous to our target were identified through BLAST+ (Camacho, et al., 2009), processed with Python and stored in a PostgreSQL database. The parameters for the BLAST search were set to retrieve proteins with a percent identity greater than 20%, an expected value less than 10^-3^, a query coverage of greater than 75%, and the total number of BLAST results limited to 50. These criteria ensured that a wide enough variety of proteins with shared similarities to the target could be found.

Because of the large number of BLAST results, a refined scoring system was needed to prioritize proteins that were both structurally and functionally similar to the target. To achieve this ranking, we developed a composite similarity score that combined sequence similarity and taxonomic proximity.

The first component of the similarity score was the percent identity (*pident*) from BLAST, normalized to a value between 0 and 1: 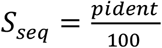

The second component was calculated based on the taxonomic distance between the BLAST target (query) and result (subject) organisms. We first queried the UniProt API (The UniProt Consortium, 2024) to find the result organism’s taxonomic ID and lineage, then searched a taxonomy tree stored inside a Neo4J graph database of related cellular organisms to capture evolutionary similarity.

We calculated the shortest path between the target organism and the result organism in the taxonomy tree, applying penalties by rank for each traversed node, further assigning the highest penalty when crossing domains. If either the target or result organism was missing from the taxonomy tree, we calculated the path by going up its lineage step by step and re-running the query, subtracting accordingly from the similarity score. The taxonomic similarity score was then calculated as:

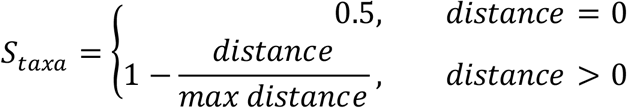

Paralogs were penalized due to their higher likelihood of having divergent or non-functional roles (Soria, et al., 2014). Specifically, if the BLAST hit was from the same organism as the target, we halved the taxonomic similarity score. The final score was the average of the sequence and taxonomic similarity components:

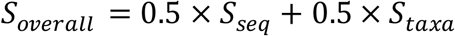

Proteins were ranked from most to least similar based on *S*_*overall*_, and hits falling below a user-defined threshold were discarded. The weighting of sequence versus taxonomic similarity can be adjusted by the user to prioritize closer evolutionary relationships or higher sequence identity. Proteins were excluded if they failed to meet the minimum similarity threshold or if the total count exceeded the user-defined limit.

### LLM Literature Mining

Using the ranked list of similar proteins from the similarity calculation module, the literature mining pipeline automatically retrieved and processed primary citations to extract purification protocols. The application queried the PDB to retrieve the primary citations of a user-defined number of most similar proteins. Proteins with inaccessible papers were discarded, while those with accessible full-text articles were downloaded from PubMed Central (PMC) in XML format and stored in memory for LLM processing.

The agents performed Python XML parsing using both XML-specific libraries and regular expressions to extract relevant metadata (e.g., title, methods section). The methods section was identified by locating the keyword ‘method’ in XML ‘<section-title>’ tags. The extracted methods section was then passed to an extraction agent, which identified and returned only the raw text related to the protein purification protocol. The extraction agent implemented error handling for malformed XML and missing sections, logging failures for user notification while continuing processing of successfully parsed articles.

All agents were developed using the PydanticAI framework (2.10.6). The extraction agent leveraged PydanticAI’s extensive data validation framework to verify the extracted purification protocol and safeguard against hallucinations, ensuring it returned only the relevant text for processing by subsequent agents. When target proteins had corresponding entries in the SSGCID database, the agents retrieved these failed protocols and provided them to subsequent agents for comparison.

All calls to LLMs in this study used Google’s Gemini-2.5-pro (Comanici, et al., 2025) (via free API access). The system, however, is model-agnostic, and users can substitute an LLM of their choice, including Open-Source models, by replacing the model specified in the PydanticAI initialization.

### Protocol Analysis Agents

To process the extracted protocol text from the literature mining pipeline, two specialized agents operated in tandem to generate different analytical outputs: a summary of mined experimental designs and an optimized purification protocol. Together, these agents enabled both an at-a-glance analysis of mined protocols and the generation of a step-by-step guide for an optimized solution. The summarization agent converted each input protocol into a standardized table with 6 columns: purification step, buffer name, buffer composition, pH, salt type, and buffer supplement, providing a quick overview of the purification protocols used for similar proteins. The summarization agent used Pydantic’s field validation to ensure output consistency, resulting in minimal variation across executions for the same input, indicating a high level of reliability. These tables, as well as the source article’s title, URL, organism, and similarity score, were then appended to a comprehensive final report. The summarization agent was also provided with the failed protocol retrieved from the SSGCID’s database and returned it as a baseline for comparison with successful approaches.

The optimizer agent compared the target’s failed protocol with the set of successful ones and generated a modified protocol based on the identified differences between the failed and successful purification attempts. Additionally, the agent was provided with structural and functional annotations, such as signal peptides and transmembrane domains, retrieved from the UniProt Knowledgebase. This allowed the agent to identify solubility hazards and incompatibilities prior to generating the protocol. Prompts were iteratively refined through testing and were made increasingly detailed as it was discovered the LLM tended to perform better when instructions were extremely clear. The optimizer agent implemented structured output formatting to ensure consistency and included confidence indicators when suggesting modifications based on limited literature evidence. This optimization analysis was then appended to the final report, creating a detailed resource that consolidated all relevant information for purifying challenging proteins.

### Web Tool

We developed a web interface using FastAPI and Svelte to provide researchers with an intuitive environment to interact with the agents. The application separates the computational workload from the user interface, using asynchronous background workers to execute the LLM pipeline. Users can input FASTA sequences or target IDs (SSGCID), which the system automatically parses to fetch relevant internal or external data. The application integrates with the SSGCID internal database to retrieve failed protocols when an SSGCID ID is detected. The interface consists of three distinct views: an input form, a live processing dashboard, and a comprehensive results report. The processing dashboard uses HTTP polling to retrieve status updates from the backend, guiding the user from initial analysis to the final generated protocol.

## Results

Starting with a given failed protocol, the multi-agent system follows a coordinated process, with the complete flowchart shown in Figure 2. First, the agents use sequence similarity tools and taxonomic lineage analysis to identify similar proteins in the Protein Data Bank (PDB) and retrieve their associated primary citations. Proteins with sufficient similarity scores and accessible literature serve as candidates for protocol rescue. Next, the extraction agent mines methodological details from the retrieved publications and extracts the raw purification protocol. This text is then analyzed by the summarizer agent, which summarizes each purification step in a table. Finally, if a failed protocol is provided, the optimizer agent compares successful protocols with the failed one to identify critical differences, proposes a revised protocol, and generates a structured report that outlines discrepancies and suggests specific protocol modifications.

**Figure 2:**
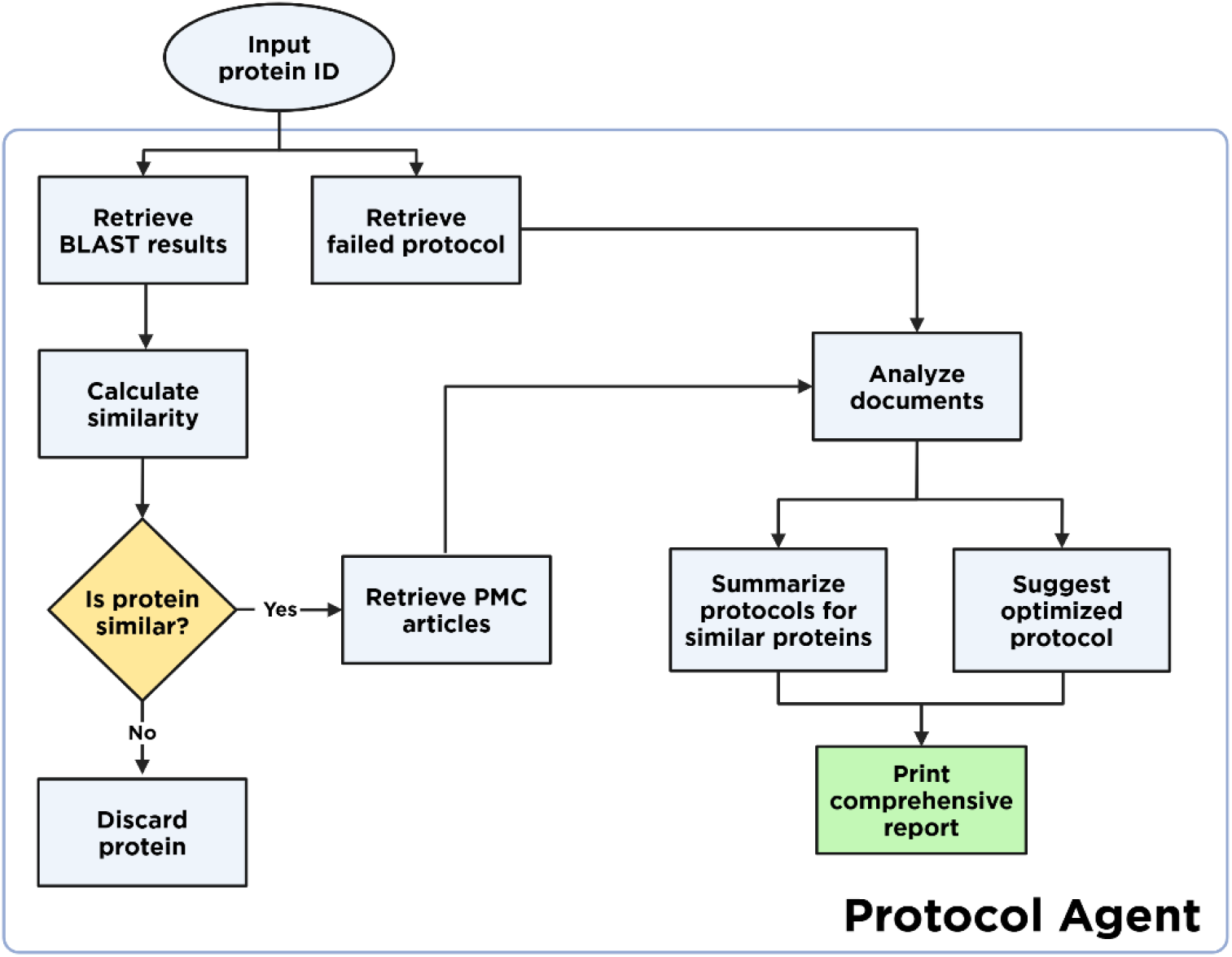
A system flowchart of the application’s process.

After developing the multi-agent system, we evaluated its performance in automating protein purification rescue workflows, assessing both the accuracy of literature extraction and the quality of generated protocol recommendations. We tested the system using a dataset of failed purification targets to demonstrate real-world applicability.

### Dataset Performance and Literature Accessibility

Our evaluation dataset consisted of failed purification cases throughout the SSGCID pipeline (Bryan, et al., 2011). We initially focused on *Mycobacterium tuberculosis* (MTB) proteins but expanded to all *Mycobacterium* proteins that expressed in soluble form, but failed large-scale purification, achieving sufficient diversity for comprehensive system testing. From an initial pool of 104 potential targets (86 non-TB mycobacterial (NTM) proteins and 18 MTB proteins), we excluded those returning no BLAST results with an expected value of less than 10^-3^ and a query coverage of greater than 75%, as well as those without legal programmatic access to PMC articles describing their purification. This resulted in a final total of 48 targets (42 NTM and 6 MTB). A primary limitation in target processing was literature accessibility: 50% of initial targets were excluded due to inaccessible PubMed Central articles, with primary citations either unpublished, lacking open access permissions, or absent entirely (Figure 3). Despite these constraints, the final targets represented a diverse selection of proteins, ranging from 1 to 463 BLAST results and up to 210 programmatically accessible PMC articles. For detailed system evaluation and case study analysis, three representative targets were selected and processed as FASTA inputs to demonstrate the complete workflow.

**Figure 3:**
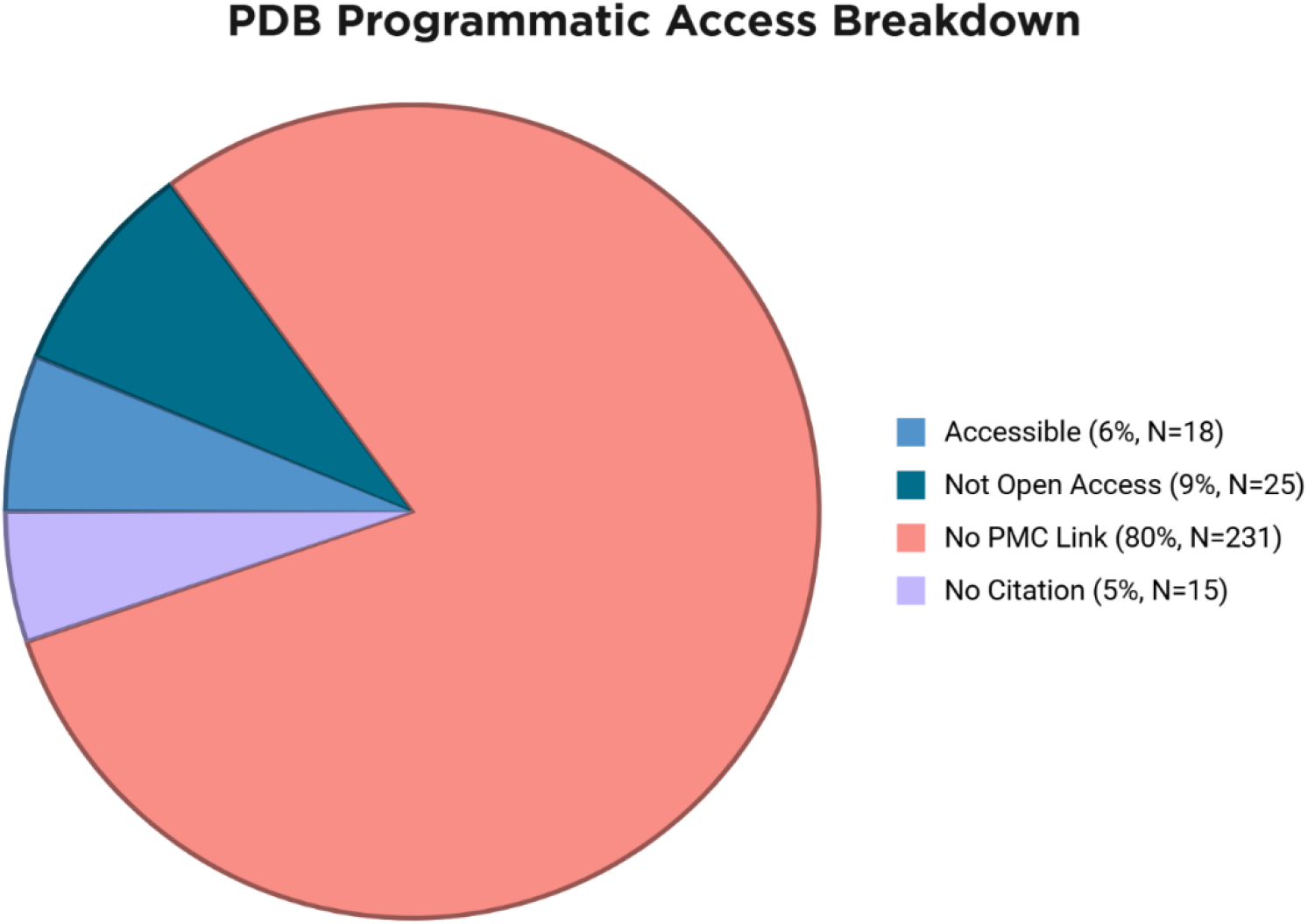
Number of inaccessible PubMed Central articles.

### Protocol Extraction and Summarization Accuracy

Through extensive prompt and context engineering, the agent produced both a detailed optimized purification protocol and a comparative analysis highlighting key differences between the failed and successful approaches. Overall, the application displayed a high level of accuracy and knowledge. A review of automated summaries by laboratory scientists experienced in protein purification found no errors in translating mined experimental details to a tabular summary. The summaries were comprehensive, and no information was lost (Table 1).

**Table 1:** Example automated purification protocol summary.

| Purification Step | Buffer Name | Buffer Composition | pH | Salt Type | Buffer Supplement |
| --- | --- | --- | --- | --- | --- |
| Lysis | lysis buffer | 20 mM HEPES, 300 mM NaCl, 5% glycerol, 30 mM Imidazole, 0.5% CHAPS, 10 mM MgCl2, 3 mM β-mercaptoethanol, 1.3 mg/ml protease inhibitor cocktail (Roche, Basel, Switzerland), and 0.05 mg/ml lysozyme | 7.4 | NaCl | 1.3 mg/ml protease inhibitor cocktail (Roche, Basel, Switzerland), and 0.05 mg/ml lysozyme |
| HisTrap FF 5 ml column - Equilibration/Binding | binding buffer | 25 mM HEPES, 300 mM NaCl, 5% Glycerol, 30 mM Imidazole, 1 mM DTT | 7.0 | NaCl | None |
| HisTrap FF 5 ml column - Elution | None | 25 mM HEPES, 300 mM NaCl, 5% Glycerol, 500 mM Imidazole, 1 mM DTT | 7.0 | NaCl | None |
| Cleavage - Dialysis | cleavage buffer | 20 mM HEPES, 500 mM NaCl, 5% Glycerol, 1 mM TCEP | 7.6 | NaCl | None |
| Ni Sepharose 6 Fast Flow resin - Wash | wash buffer | 20 mM HEPES, 300 mM NaCl, 40 mM imidazole, 1 mM TCEP 5% glycerol | 7.0 | NaCl | None |
| Superdex 75 26/60 column - Equilibration | SEC buffer | 20 mM HEPES, 300 mM NaCl, 5% glycerol and 1 mM TCEP | 7.0 | NaCl | None |

To further validate system performance, we conducted detailed analysis of three representative test cases, which demonstrated the system’s ability to generate coherent, scientifically sound optimization suggestions. The optimizer agent consistently identified meaningful differences between failed and successful protocols, focusing on critical parameters such as high uses of abrasive chemicals like imidazole, low centrifugation speeds at 10,000 RPM, etc. Generated protocols exhibited logical consistency with established protein purification principles and included step-by-step procedures that expert reviewers deemed both promising and viable in standard laboratory settings.

### Web Interface

The Django web interface served as both the user entry point and the coordination hub for the entire pipeline. Users could enter a protein’s FASTA, PDB, or SSGCID ID, and upload their own failed protein purification protocol if no SSGCID protocol was available. The web tool automatically detected the input type and responded accordingly. If an SSGCID ID was provided, it queried the SSGCID database. Otherwise, it skipped this step and performed the BLAST search. During processing, users were directed to a loading screen that gave periodic updates of the agents’ progress, allowing them to monitor progress and understand what the application was doing. Once the agents finished running in accordance with the previous sections, users were redirected to the results page that displayed the outputs of the summarization and optimizer agents in a structured format with downloadable reports and links to source literature.

## Discussion

### Potential of LLMs in a Lab Setting

The development, implementation, and testing of the multi-agent LLM system demonstrate that it successfully automates the labor-intensive process of protein purification rescue, reducing a typically hours-long manual workflow to a two-minute automated analysis. Beyond improvements in efficiency, the system’s accuracy in mining protocols and generating scientifically sound optimization recommendations show the potential for specialized LLM applications to overcome bottlenecks in laboratory research workflows.

The test of protocol summarization reveals no errors in how the system extracted and organized experimental details from the literature. This accuracy shows the effectiveness of LLM mining of scientific literature, especially when leveraging PydanticAI’s validation framework to prevent hallucinations. One encouraging aspect of the agents’ behavior is their appropriate handling of ambiguity; they flag uncertainties and present options rather than guessing. This approach is essential in laboratory settings, where researchers require a full spectrum of potential choices rather than automated selections made by an LLM.

Generated optimized protocols were coherent and demonstrated an accurate knowledge of physical chemistry and structural genomics. According to experts during their review, these optimized protocols often suggested changes that a scientist in the lab would be likely to consider when asked to rescue a failed protocol. The optimizer agent consistently identified meaningful protocol differences, suggesting that LLMs can replicate expert analysis when provided with the appropriate prompts, grounding material, and validation frameworks.

### Limitations

A fundamental limitation of this retrieval-based system is that its output is only as good as the protocols it extracts from literature. While LLMs excel at summarization and analysis tasks, they struggle to generate novel approaches, especially in a laboratory setting (Bender, et al., 2021). At its core, this application replicates the process by which scientists typically rescue failed protein purifications and may struggle to suggest innovative protocols, especially when the target protein has few closely related proteins in the existing literature. Future directions could incorporate more biochemical or physics-based knowledge to enable novel design generation while maintaining experimental grounding. Additionally, integration with protein structure prediction tools or other bioinformatics tools could help the system expand beyond documented approaches (Jumper, et al., 2021).

Another limitation is literature accessibility, with the vast majority of PDB primary citations unavailable for several reasons (Figure 3). The most common limiting factor is that the citation in the PDB is not on PubMed Central. Additionally, some PDB entries lack a primary citation at all, as the corresponding paper has not or will not be published. Even when the PMC article exists, it might lack open access permissions for programmatic full-text retrieval. This bottleneck highlights a challenge with LLM-based automation: the reliance upon limited open-access scientific literature for automated analysis.

### Advantages of the Multi-Agent Architecture

The modular design of specialized agents provides significant advantages in our implementation. Each agent can be independently optimized, validated, and updated without changing other parts of the pipeline. In practice, this architecture offers flexibility for workflow modifications. Each agent functions as a self-contained component that can operate so long as it is given a viable input. Therefore, new agents can be inserted to alter or oversee any part of the pipeline. Some future agents that could be added include an LLM judge to act as an online evaluation framework and prevent hallucinations, fact-checking each output and using tools or stored knowledge to prevent factual errors. Another agent could offer physical chemistry and genomics knowledge, allowing the application to generate novel protocol steps based on its expertise.

Other ways this application could be expanded are to include downstream steps of the protein purification pipeline, such as crystallography. The workflows for these processes require similar knowledge and abilities, such that the application could easily expand to encompass them.

In addition to automating a time-consuming phase of protein purification, this application provides a proof of concept for LLMs and agentic architecture in a laboratory context. Although not yet widely adopted, these technologies have the potential to transform research workflows. By reducing the burden of repetitive tasks, they allow for more efficient research and accelerated discovery. The success of this specialized LLM application supports the broader adoption of these technologies, so long as careful attention is paid to scientifically sound grounding, the prevention of hallucinations, and integration with existing research tools.

## Code Availability

The source code for this tool may be found in the GitHub repository (https://github.com/jeffery-ye/agentic-protein-purification) or archived through Zenodo (https://doi.org/10.5281/zenodo.18852283).

## Acknowledgements

This project has been funded in whole with Federal funds from the National Institute of Allergy and Infectious Diseases (NIAID), NIH, Department of Health and Human Services, under Contract No. 75N93022C00036.

